# Exogenous phosphatidic acid reduces acetaminophen-induced liver injury in mice by activating hepatic interleukin-6 signaling through inter-organ crosstalk

**DOI:** 10.1101/2020.12.22.423312

**Authors:** Melissa M. Clemens, Stefanie Kennon-McGill, Joel H. Vazquez, Owen W. Stephens, Erich A. Peterson, Donald J. Johann, Felicia D. Allard, Eric U. Yee, Sandra S. McCullough, Laura P. James, Brian N. Finck, Mitchell R. McGill

## Abstract

We previously demonstrated that endogenous phosphatidic acid (PA) promotes liver regeneration after acetaminophen (APAP) hepatotoxicity in mice. Based on that, we hypothesized that exogenous PA is also beneficial. To test that, we treated mice with a toxic APAP dose at 0 h, followed by PA or vehicle at multiple timepoints. We then collected blood and liver at 6, 24, and 52 h. Post-treatment with PA protected against liver injury at 6 h, and the combination of PA and N-acetyl-cysteine (NAC) further reduced injury compared to NAC alone. Interestingly, PA had no effect on major early mechanisms of APAP toxicity, including APAP bioactivation, oxidative stress, JNK activation, and mitochondrial damage. However, transcriptomics revealed that PA activated interleukin-6 (IL-6) signaling in the liver, and IL-6 was increased in serum from PA-treated mice. Furthermore, PA did not protect against APAP in IL-6-deficient mice. In addition, IL-6 expression increased 18-fold in adipose tissue after PA, indicating that adipose tissue is a likely source of the increased IL-6 due to PA treatment. Surprisingly, however, exogenous PA did not alter regeneration, despite the widely accepted role of IL-6 in liver repair. These data reinforce the protective role of IL-6 in APAP hepatotoxicity, provide new insight into the role of IL-6 in liver regeneration, and indicate that exogenous PA or PA derivatives may one day be a useful adjunct treatment for APAP overdose with NAC.

## 1. Introduction

Acetaminophen (APAP) is a popular analgesic and antipyretic drug (Kaufman et al., 2002), but overdose causes severe acute liver injury. It is currently the leading cause of acute liver failure (ALF) throughout much of the world (Lee, 2008). Conversion of APAP to the reactive metabolite N-acetyl-*p*-benzo quinoneimine (NAPQI) initiates the hepatotoxicity. NAPQI binds to free sulfhydryl groups on amino acid residues, depleting hepatic glutathione and damaging proteins (Jollow et al., 1973; Mitchell et al., 1973; McGill and Hinson, 2020). The protein binding leads to mitochondrial dysfunction and oxidative stress (Jaeschke, 1990; Cover et al., 2005), which activates the c-Jun N-terminal kinases 1/2 (JNK) and other kinases (Gunawan et al., 2006; Hanawa et al., 2008; Nakagawa et al., 2008; Ramachandran et al., 2013). Activated JNK translocates from the cytosol to mitochondria, where it exacerbates the mitochondrial dysfunction by reducing mitochondrial respiration (Hanawa et al., 2008; Win et al., 2016). Eventually, the mitochondrial permeability transition occurs (Kon et al., 2004; Reid et al., 2005), and the mitochondrial damage causes release of endonucleases from mitochondria, which then cleave nuclear DNA (Bajt et al., 2006). The affected hepatocytes die by necrosis (Gujral et al., 2002; McGill etal., 2011; 2012).

Phosphatidic acid (PA) is a critically important lipid in all prokaryotic and eukaryotic cells. It is the simplest diacylated glycerophospholipid, having a bare phosphate head group. In cell and organelle membranes, the small size and negative charge of the head group likely promotes negative curvature that may be important for membrane fission (Kooijman et al., 2003). It is also a major metabolic intermediate, serving as a key precursor for synthesis of all other phospholipids as well as triglycerides (Pokotyla et al., 2018). Finally, it is a major lipid second messenger that is known to play a role in nutrient sensing and cell proliferation via mechanistic target of rapamycin (mTOR) signaling (Fang et al., 2001; Foster, 2013; Pokotyla et al., 2018).

We recently demonstrated that phosphatidic acid (PA) is beneficial after APAP-induced liver injury in mice through an entirely novel mechanism (Lutkewitte et al., 2018; Clemens et al., 2019a). Briefly, we found that *endogenous* PA is elevated in the liver after APAP overdose, and that it promotes cell proliferation and therefore liver repair by regulating glycogen synthase kinase 3β (GSK3β) (Lutkewitte et al., 2018; Clemens et al., 2019). However, we did not test the effect of *exogenous* administration of PA on APAP-induced liver injury. In the present study, we hypothesized that exogenous PA is beneficial in a mouse model of APAP overdose. Our data demonstrate that it reduces APAP hepatotoxicity by increasing systemic interleukin-6 (IL-6) from adipose tissue, which in turn upregulates protective IL-6 signaling in the liver.

## 2. Experimental procedures

### 2.1. Animals

Male wild-type (WT) C57BL/6J mice and IL-6 knockout mice (IL-6 KO; B6.129S2-Il6^tm1Kopf^/J) between the ages of 8 and 12 weeks were obtained from The Jackson Laboratory (Bar Harbor, ME, USA). The mice were housed in a temperature-controlled 12 h light/dark cycle room and allowed free access to food and water. The APAP and PA solutions were prepared fresh on the morning of each experiment. APAP was prepared by dissolving 15 mg/mL APAP (Sigma, St. Louis, MO, USA) in 1x PBS with gentle heating and intermittent vortexing. The PA solution was prepared by re-constituting purified egg PA extract (Avanti Polar Lipids, Alabaster, AL, USA) at 10 mg/mL in 10% DMSO in 1x PBS, warming to 80°C for 20-30 min with intermittent mixing to obtain a uniform hazy suspension, and cooling to room temperature immediately before injection. To determine if PA affects liver injury, WT mice (n = 5-10 per group) were fasted overnight then injected (i.p.) with 250 mg/kg APAP at 0 h, followed by 10% DMSO vehicle (Veh) or 20 mg/kg PA (i.p.) at 2 h. Blood and liver tissue were collected at 6 h. We chose the 20 mg/kg dose of PA because it is commonly recommended when taken as a dietary supplement in humans. To determine if the combination of N-acetylcysteine (NAC) and PA reduces injury compared to NAC alone, some mice were injected with APAP at 0 h followed by 300 mg/kg NAC (dissolved in 1x PBS) and either PA or vehicle at 2 h (n = 7 per group). Blood was collected at 6 h. We chose the 300 mg/kg dose of NAC because it is approximately 2-fold greater than the typical loading dose in humans after APAP overdose. Using this high dose of NAC ensures that our results comparing NAC with APAP+NAC are conservative and robust. For transcriptomics, the original PA experiment was repeated at the 6 h time point with addition of a vehicle-only control group (n = 5 per group). To determine if PA protection depends upon IL-6, the experiment was repeated again at the 6 h time point using IL-6 KO mice (n = 5-6 per group) and a similar but higher dose of APAP (350 mg/kg). The change in APAP dose in the latter experiment was due to an adjustment made to our university animal use protocol during the course of the study and unrelated to our data from these experiments. Finally, to test the role of Kupffer cells, the original PA experiment was repeated at the 6 h time point with WT mice (n = 10 per group) after 24 h i.v. (tail vein) pre-treatment with 0.2 mL of 17 mM liposomal clodronate (Clodrosome, Brentwood, TN, USA). All study protocols were approved by the Institutional Animal Care and Use Committee of the University of Arkansas for Medical Sciences.

### 2.2. Subcellular fractionation

Right and caudate liver lobes were homogenized in ice cold isolation buffer containing 220 mM mannitol, 70 mM sucrose, 2.5 mM HEPES, 10 mM EDTA, 1mM ethylene glycol tetra-acetic acid and 0.1% bovine serum albumin (pH 7.4) using a Thermo-Fisher Bead Mill. Subcellular fractions were obtained by differential centrifugation. Samples were centrifuged at 2,500 x g for 10 min to blood cells and debris. Supernatants were then centrifuged at 20,000 x g for 10 min to pellet mitochondria. The supernatant was retained as the cytosol fraction. Pellets containing mitochondria were then resuspended in 100 μL of isolation buffer and freeze-thawed three times using liquid nitrogen to disrupt the mitochondrial membranes. Protein concentration was measured in both the mitochondrial and cytosol fractions using the BCA assay, and the samples were used for western blotting as described below.

### 2.3. Clinical Chemistry

Alanine aminotransferase (ALT) was measured in serum using a kit from Point Scientific Inc. (Canton, MI, USA) according to the manufacturer’s instructions.

### 2.4. Histology

Liver tissue sections were fixed in 10% formalin. For hematoxylin & eosin (H&E) staining, fixed tissues were embedded in paraffin wax, then 5 μm sections were mounted on glass slides and stained according to a standard protocol. Necrosis was quantified in the H&E-stained sections by two independent, fellowship-trained, hepatobiliary pathologists who were both blinded to sample identity. Percent necrosis was then averaged for each animal. For Oil Red O staining, fixed tissues were embedded in OCT compound and rapidly frozen by placing on a metal dish floating in liquid nitrogen. 8 μm sections were cut and mounted on positively-charged glass slides. The sections were allowed to dry for 30 min at room temperature, then treated with 60% isopropanol for 5 min, followed by freshly prepared Oil Red O solution in isopropanol for 10 min, and then 60% isopropanol for an additional 2 min. The sections were then rinsed with PBS, treated with Richard-Allan Gill 2 hematoxylin solution (Thermo Fisher, Waltham, MA, USA) for 1 min, and rinsed again with PBS before cover-slipping. Digital images were taken using a Labomed Lx400 microscope with digital camera (Labo American Inc., Fremont, CA, USA).

### 2.5. Western Blotting

Liver tissues were homogenized in 25 mM HEPES buffer with 5 mM EDTA, 0.1% CHAPS, and protease inhibitors (pH 7.4). Protein concentration was measured using a bicinchoninic acid (BCA) assay. The samples were then diluted in homogenization buffer, mixed with reduced Laemmli buffer, and boiled for 1 min. Equal amounts (60 μg protein) were added to each lane of a 4-20% Tris-glycine gel. After electrophoresis, proteins were transferred to PVDF membranes and blocked with 5% milk in Tris-buffered saline with 0.1% Tween 20. Primary monoclonal antibodies were purchased from Cell Signaling Technology (Danvers, MA, USA): p-JNK (Cat. No. 4668), JNK (Cat. No. 9252), AIF (Cat. No. 5318), Cytochrome-C (Cat. No.11940), AIF (Cat. No. 5318), GSK3β (Cat. No. 9315), phospho-GSK3β (Cat. No. 9323), and β-actin (Cat. No. 4967). All primary antibodies were used at 1:1000 dilution. Secondary antibodies were purchased from LiCor Biosciences (Lincoln, NE, USA): IRDye 680 goat anti-mouse IgG (Cat. No. 926-68070) and IRDye 800CW goat anti-rabbit IgG (Cat. No. 926-32211). All secondary antibodies were used at 1:10,000 dilution. Bands were visualized using the Odyssey Imaging System (LiCor Biosciences, Lincoln, NE, USA).

### 2.6. Glutathione measurement

Total glutathione (GSH+GSSG) and oxidized glutathione (GSSG) were measured using a modified Tietze assay, as we previously described in detail (McGill and Jaeschke, 2015).

### 2.7. APAP-protein adduct measurement

APAP-protein adducts were measured using high pressure liquid chromatography (HPLC) with electrochemical detection, as previously described (Muldrew et al., 2002; McGill et al., 2013).

### 2.8. Transcriptomics

The Supporting Information section contains all details concerning RNA-seq sample prep, next generation sequencing, and bioinformatic analyses.

### 2.9. Statistics

Normality was assessed using the Shapiro-Wilk test. Normally distributed data were analyzed using a t-test for comparison of two groups or one-way ANOVA with post-hoc Student-Neuman-Keul’s for comparison of three or more groups. Data that were not normally distributed were analyzed using a nonparametric Mann-Whitney U test for comparison of two groups, or one-way ANOVA on ranks with post-hoc Dunnet’s test to compare three or more. All statistical tests were performed using SigmaPlot 12.5 software (Systat, San Jose, CA, USA).

## 3. Results

### 3.1. Exogenous PA reduces liver injury at 6 h after APAP overdose

To determine the effect of exogenous PA treatment on APAP-induced liver injury, we treated mice with APAP at 0 h followed by PA or vehicle at 2 h. We then collected blood and liver tissue at 6 h. We observed a significant reduction in serum ALT values in the PA-treated mice at 6 h post-APAP (Fig. 1A). Two blinded, fellowship-trained hepatobiliary and GI pathologists independently confirmed the reduction in injury based on histology (Fig. 1B and Table 1). NAC is the current standard-of-care treatment for APAP-induced liver injury in patients. To determine if the combination of PA and NAC can further reduce injury after APAP overdose compared to the standard-of-care alone, we treated mice with APAP followed by 300 mg/kg NAC and either vehicle or PA. The combination of NAC plus PA significantly decreased serum ALT compared to NAC plus vehicle (Table 2). Together, these data demonstrate that exogenous PA can reduce liver APAP-induced liver injury and indicate that it could be useful as an adjunct treatment for APAP overdose.

**Table 1.** Percent necrosis in H&E-stained liver sections.

| **Percent**<br>**necrosis:** | **0-10%** | **11-25%** | **26-50%** | **>50%** |
| --- | --- | --- | --- | --- |
| **Vehicle** | 20% | 60% | 20% | 0 |
| **PA** | 100% | 0 | 0 | 0 |

**Table 2.** Comparison of NAC/Veh and NAC/PA after APAP overdose.

| **Treatment** | **ALT (U/L), mean±SE** | **p-value** |
| --- | --- | --- |
| **APAP+NAC+Veh** | 170±31 |  |
| **APAP+NAC+PA** | 69±12 | 0.011 |

**Figure 1.**
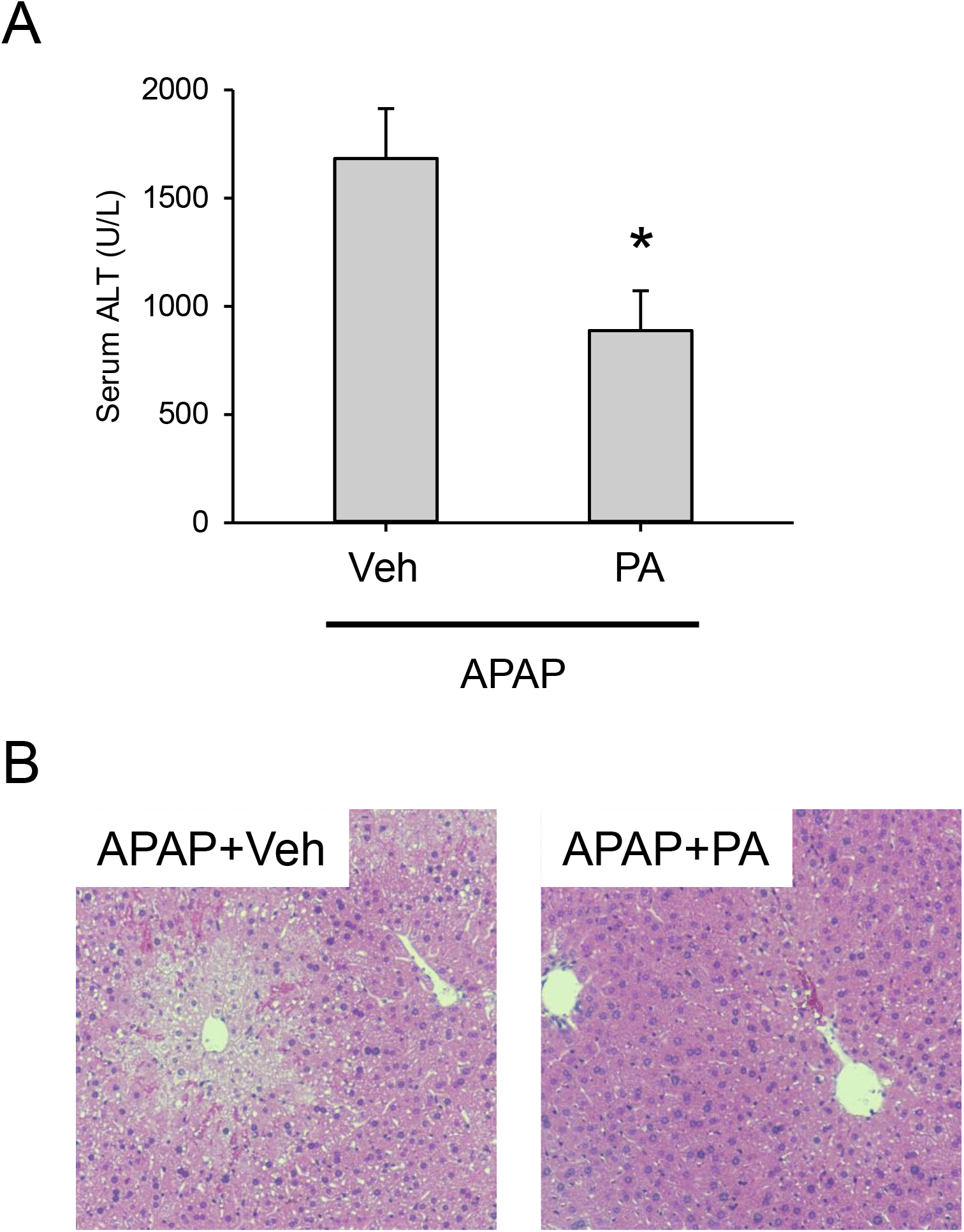
Post-treatment with exogenous PA protects against early APAP hepatotoxicity. Mice were treated with 250 mg/kg APAP at 0 h, followed by vehicle (Veh) or PA at 2 h. Blood and liver tissue were collected at 6 h. (A) Serum ALT activity. (B) H&E-stained liver sections. Data expressed as mean ± SE for n = 10 per group. *p<0.05 vs. APAP plus Veh.

### 3.2. Exogenous PA does not affect the canonical mechanisms of APAP-induced liver injury

Next, we sought to determine the mechanisms by which exogenous PA reduces early APAP hepatotoxicity. The initiating step in APAP-induced liver injury is formation of the reactive metabolite N-acetyl-*p*-benzoquinone imine (NAPQI), which depletes glutathione and binds to proteins. To determine if the decrease in liver injury at 6 h was due to an effect on NAPQI formation, we measured total glutathione (GSH + GSSG) and APAP-protein adducts in the liver. We did not detect a significant difference between the APAP plus vehicle and APAP plus PA groups in either parameter (Fig. 2A,B).

**Figure 2.**
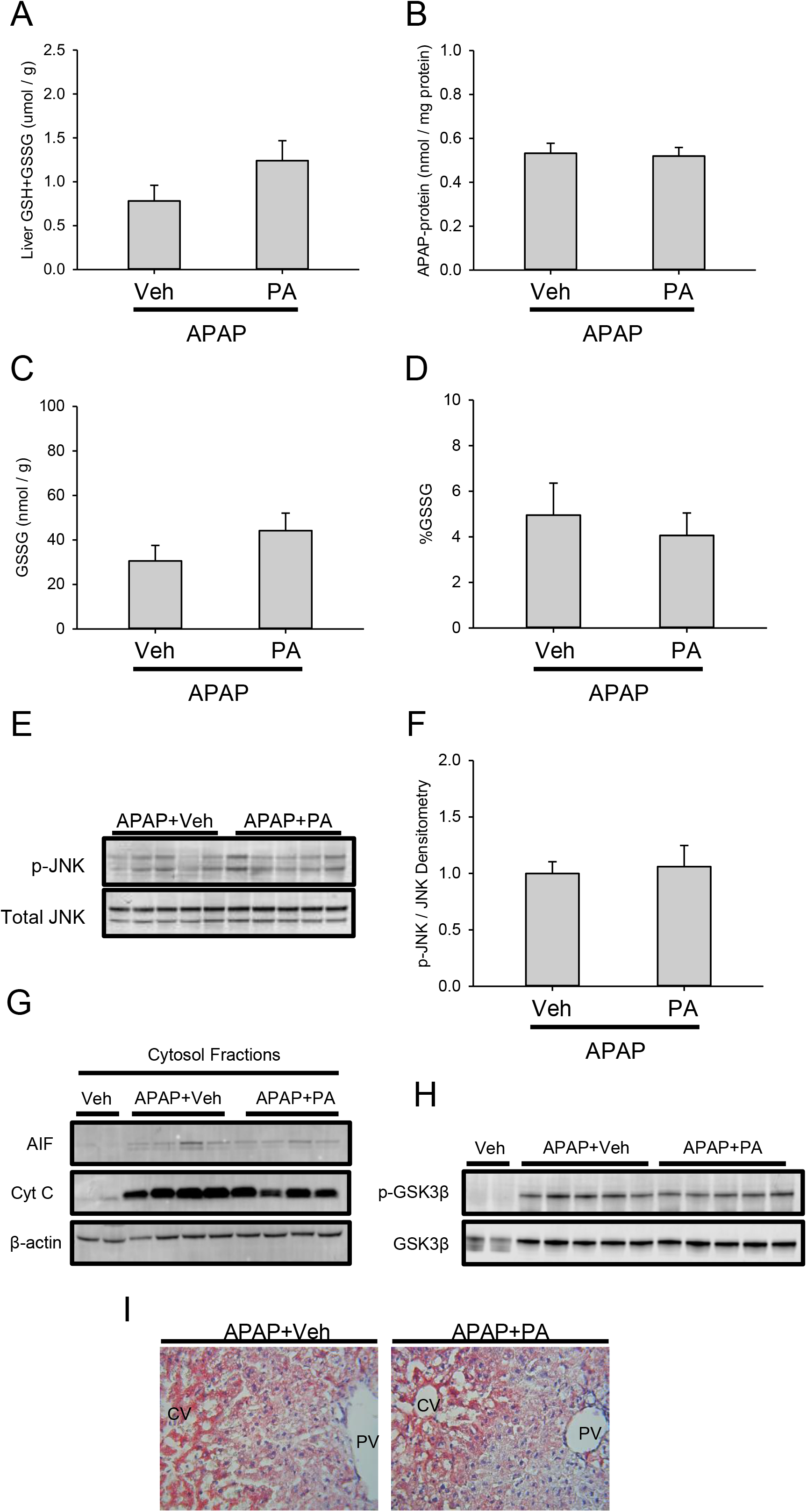
Post-treatment with exogenous PA does not affect canonical mechanisms of APAP hepatotoxicity. Mice were treated with 250 mg/kg APAP at 0 h, followed by vehicle (Veh) or PA at 2 h. Liver tissue was collected at 6 h. (A) Total glutathione (GSH+GSSG) in liver. (B) APAP-protein adducts in liver. (C) Absolute oxidized glutathione (GSSG) in liver. (D) GSSG as the percentage of total glutathione (%GSSG). (E) Immunoblots for total and phosphorylated JNK. (F) JNK densitometry. (G) Immunoblots for AIF, cytochrome c, and β-actin. (H) Immunoblots for total and phosphorylated (Ser9) GSK3β. (I) Oil Red O staining in liver sections. Data expressed as mean ± SE for n = 5 per group. No statistically significant differences were detected.

To determine if PA protects by preventing the early mitochondrial dysfunction and oxidative stress after APAP overdose, we measured GSSG in the liver. There was no significant difference in either total GSSG or the percentage of glutathione in the form of GSSG (%GSSG) between the two groups (Fig. 2C,D). To test if JNK activation and/or mitochondrial translocation were affected, we immunoblotted for phosphorylated and total JNK. Again, we could not detect a difference between the groups (Fig. 2E,F). To determine if PA had an effect on mitochondrial damage downstream of JNK, and therefore mitochondrial rupture, we also immunoblotted for AIF and cytochrome c in cytosolic fractions, and again no differences were detected (Fig. 2G). Because we previously found that endogenous PA can regulate GSK3β activity through Ser9 phosphorylation (Clemens et al., 2019a) and because active GSK3β is known to exacerbate APAP-induced liver injury (Shinohara et al., 2010), we also measured GSK3β Ser9 phosphorylation but once again observed no differences (Fig. 2H). Finally, as an additional indicator of mitochondrial function, we performed Oil Red O staining of triglycerides in frozen liver sections to assess lipid oxidation. Consistent with previous studies (Bhattacharyya et al., 2014; Borude et al., 2018), we observed Oil Red O accumulation in the damaged hepatocytes within centrilobular regions, indicating loss of β-oxidation due to mitochondrial damage, but again we saw no apparent difference between the groups (Fig. 2I). Altogether, these data largely rule out an effect of PA on APAP bioactivation, oxidative stress, and overt mitochondrial damage.

### 3.3. Exogenous PA protects through IL-6 signaling in the liver

To identify other mechanisms by which PA might reduce early APAP hepatotoxicity, we performed next generation RNA sequencing in liver tissue from mice treated with vehicle only, APAP plus vehicle, and APAP plus PA. We found that 6,192 genes were differentially expressed between the vehicle only and the APAP plus vehicle groups. Consistent with the protein alkylation, oxidative stress, and inflammation known to occur in APAP hepatotoxicity, gene ontology (biological processes; GO:BP) analysis revealed that genes involved in protein refolding, cell responses to chemical stimulus, and toll-like receptor signaling were increased by APAP, while various cell growth and cell signaling processes were decreased (Fig. 3A). 388 genes were differentially expressed between the APAP plus vehicle and APAP plus PA groups. This was insufficient for complete GO analysis, but it is notable that the GO:BP term “acute inflammatory response” was over-represented in the APAP plus PA group when using a log2 fold-change threshold of 1. Furthermore, hierarchical clustering analysis (Fig. 3B) showed clear separation of the APAP plus vehicle and APAP plus PA groups across the five biological replicates per group. Importantly, upstream analysis (Ingenuity Pathway Analysis [IPA]) revealed activation of signaling downstream of IL-6 and its target transcription factor signal transducer and activator of transcription 3 (Stat3) (Table 3). Recent studies have demonstrated that IL-6 is protective in APAP hepatotoxicity (Gao et al., 2019), and it was previously demonstrated that treatment with exogenous PA at doses similar to those we used here rapidly increase serum IL-6 concentration (Lim et al., 2003). Thus, to confirm that PA increased serum IL-6 in our experiment, we measured IL-6 protein in serum at 6 h post-APAP. Importantly, IL-6 was significantly elevated in the APAP plus PA mice compared to the APAP plus vehicle animals (Fig. 3C). Together, these data indicate that PA may protect against APAP toxicity by activating IL-6.

**Table 3.**
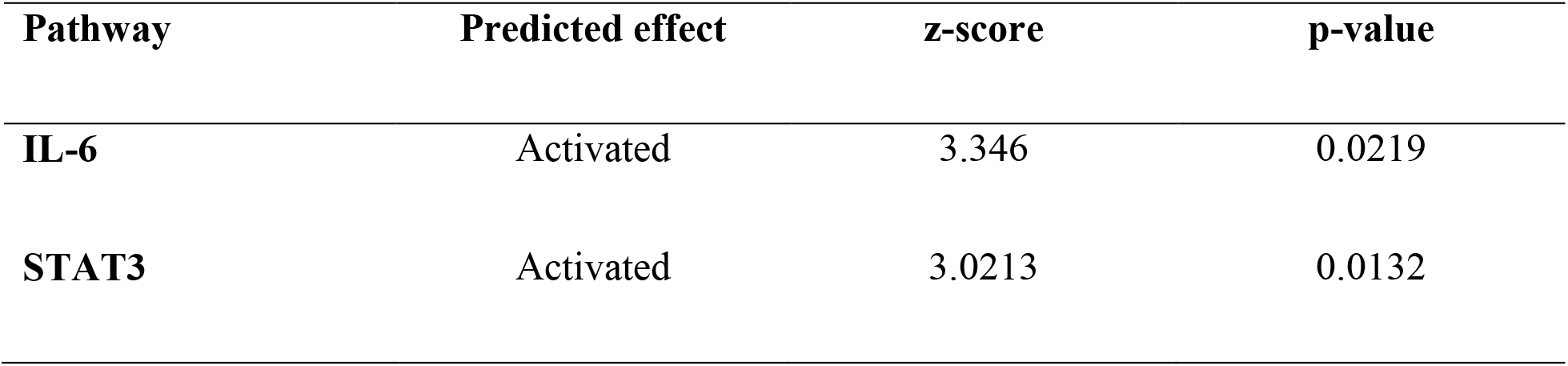
Signaling pathways affected by exogenous PA after APAP overdose.

**Figure 3.**
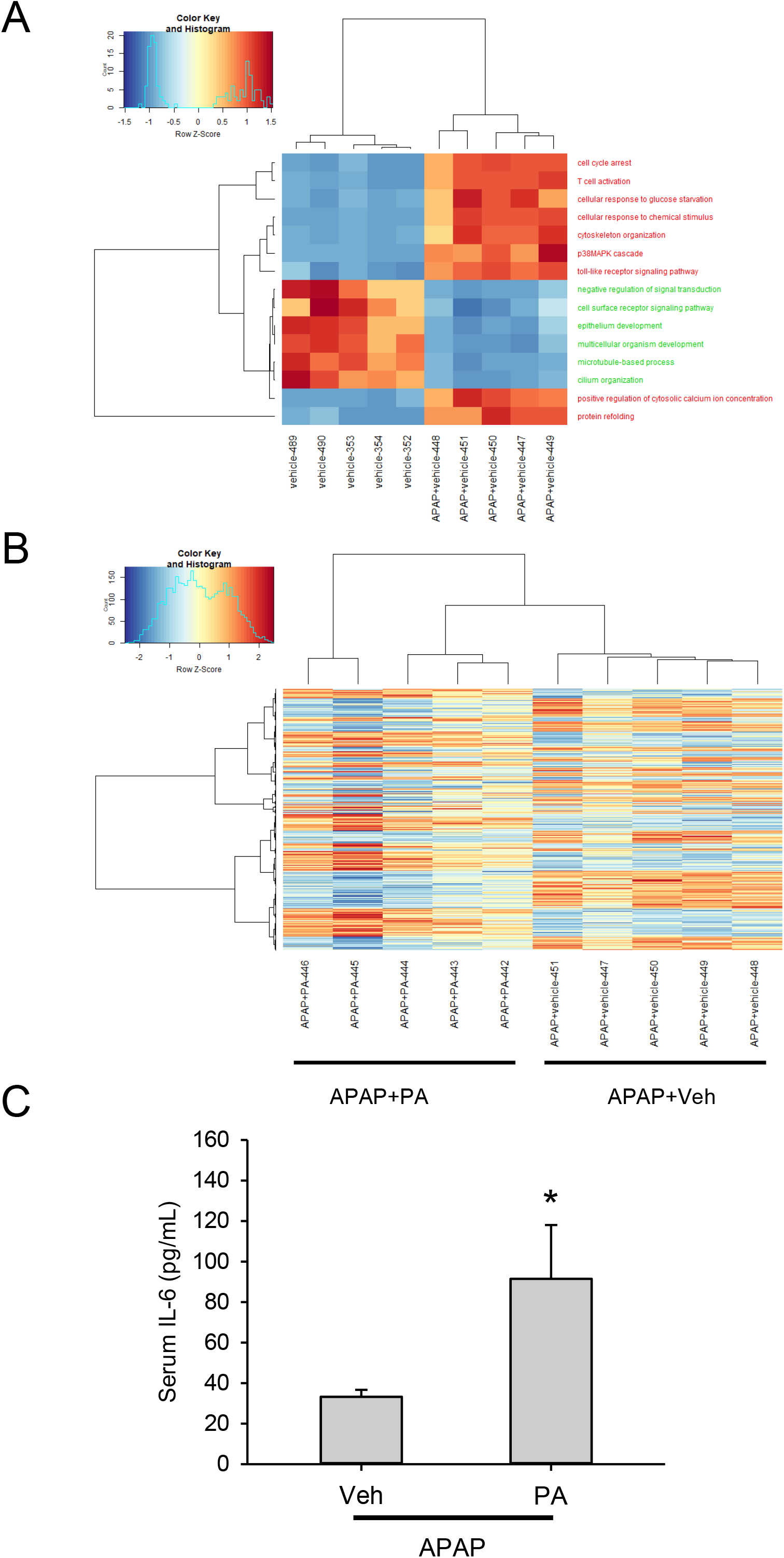
Post-treatment with exogenous PA activates IL-6 signaling in the liver. Mice were treated with 250 mg/kg APAP or vehicle alone at 0 h, followed by vehicle (Veh) or PA at 2 h. Blood and liver tissue were collected at 6 h. (A) Gene ontology (Biological Process) analysis of vehicle alone vs. APAP+vehicle. (B) Hierarchical clustering of genes in the APAP+vehicle and APAP+PA groups. (C) Serum IL-6 values. Data expressed as mean ± SE for n = 5 per group. *p<0.05 vs. APAP plus Veh.

To confirm that exogenous PA affects APAP-induced liver injury through IL-6, we compared the effect of exogenously administered PA on APAP hepatotoxicity in WT and IL-6 KO mice at 6 h post-APAP. Importantly, PA did not reduce liver injury in the KO mice, despite protecting in the WT mice in the same experiment (Fig. 4). In fact, it appeared to worsen injury in the IL-6 KO mice. These data clearly demonstrate that IL-6 is necessary for the protection provided by exogenous PA in WT mice, and support previous work indicating that IL-6 is protective in APAP-induced liver injury overall.

**Figure 4.**
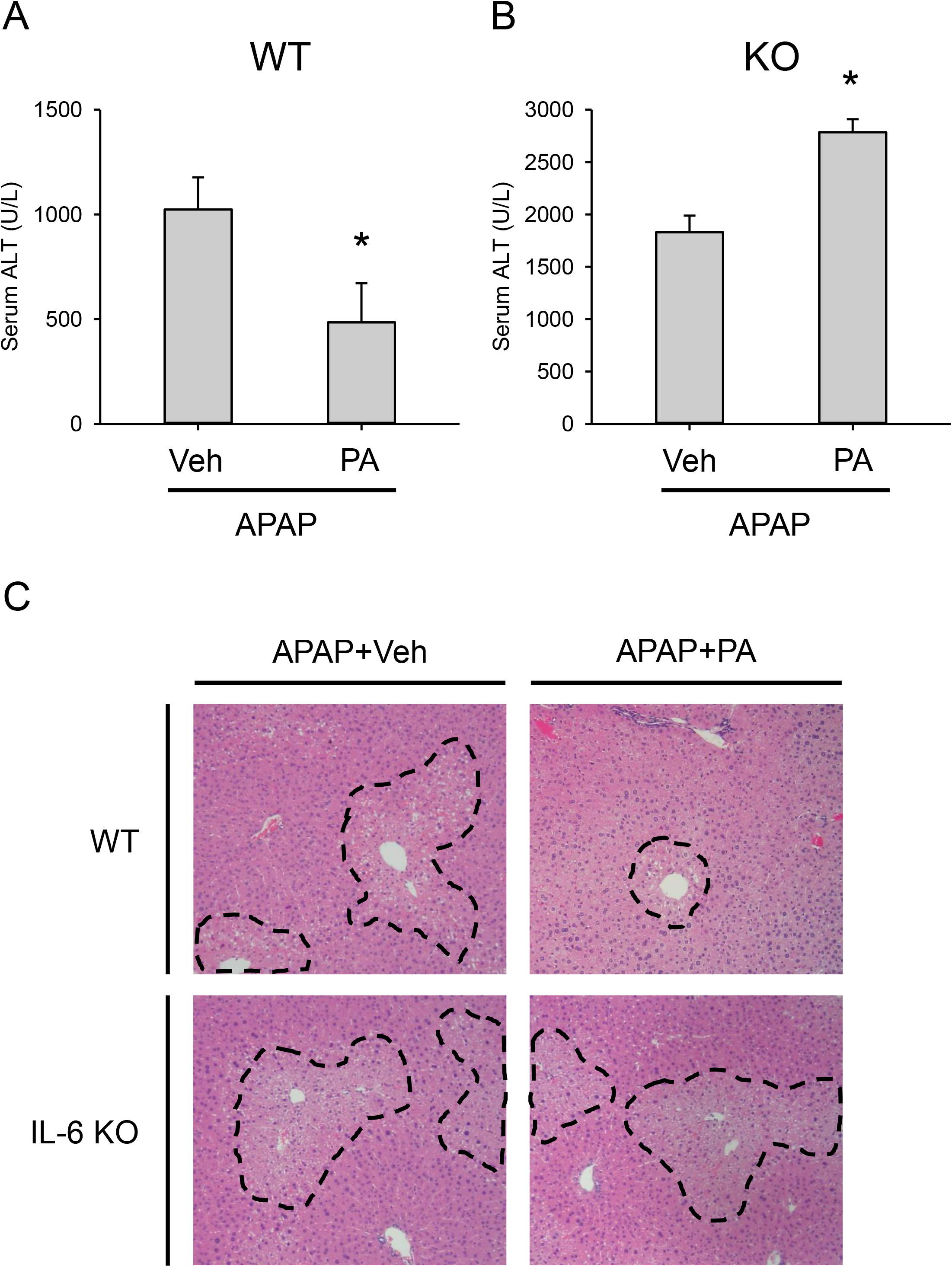
Post-treatment with exogenous PA does not protect in IL-6 KO mice. WT and IL-6 KO mice were treated with 350 mg/kg APAP at 0 h, followed by vehicle or PA at 2 h. Blood and liver tissue were collected at 6 h. (A) Serum ALT activity in WT mice. (B) Serum ALT activity in KO mice. (C) H&E-stained liver sections from both genotypes. Data expressed as mean ± SE for n = 5-6 per group. *p<0.05 vs. APAP plus Veh.

### 3.4. Adipose tissue is a likely source of increased IL-6 after PA treatment

Multiple liver cell types express IL-6, but Kupffer cells (KCs) are the major producers. To determine if the increase in IL-6 caused by treatment with exogenous PA is due to increased expression of IL-6 in KCs or other liver cells, we measured IL-6 mRNA in liver tissue in the APAP plus vehicle and APAP plus PA groups. We could not detect a significant difference in IL-6 expression between the two groups (Fig. 6A). Because KCs account for only a small portion of cells in the liver, it is possible that total liver mRNA has poor sensitivity to detect changes specifically within KCs. Thus, to further test if KCs are the source of IL-6 after PA treatment, we pre-treated mice with liposomal clodronate to ablate macrophages. The following day, we administered APAP followed by either PA or vehicle. Blood and liver tissue were collected at 6 h post-APAP. Surprisingly, serum ALT was still significantly reduced by PA (Fig. 6B), despite depletion of the liver macrophages (Fig. 6C). These data indicate that the liver itself is probably not the major source of IL-6 after PA treatment.

**Figure 5.**
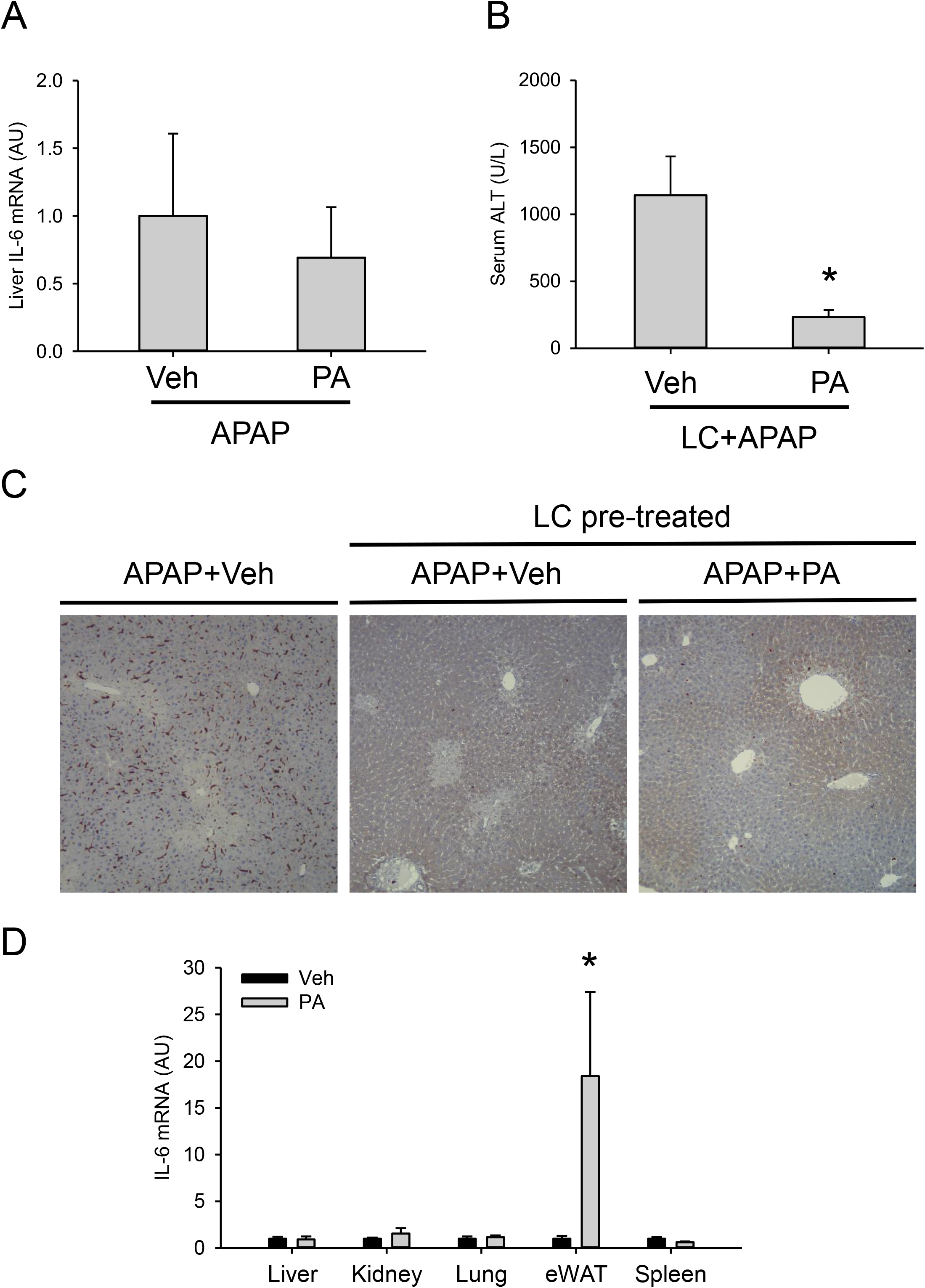
The source of IL-6 is extrahepatic and likely includes white adipose tissue. In one experiment, mice were treated with 250 mg/kg APAP at 0 h, followed by vehicle (Veh) or PA at 2 h. Where indicated, mice were pre-treated for 24 h with liposomal clodronate (LC). Blood and liver tissue were collected at 6 h. In a second experiment, mice were treated with 20 m/kg PA or vehicle and various tissues were collected 4 h later. (A) Liver IL-6 mRNA from the first experiments. (B) Serum ALT activity from the first experiment. (C) F4/80 immunohistochemistry in liver tissue sections from the first experiment. (D) IL-6 mRNA from the second experiment. Data expressed as mean ± SE for n = 5-10 per group. *p<0.05 vs. APAP plus Veh.

**Figure 6.**
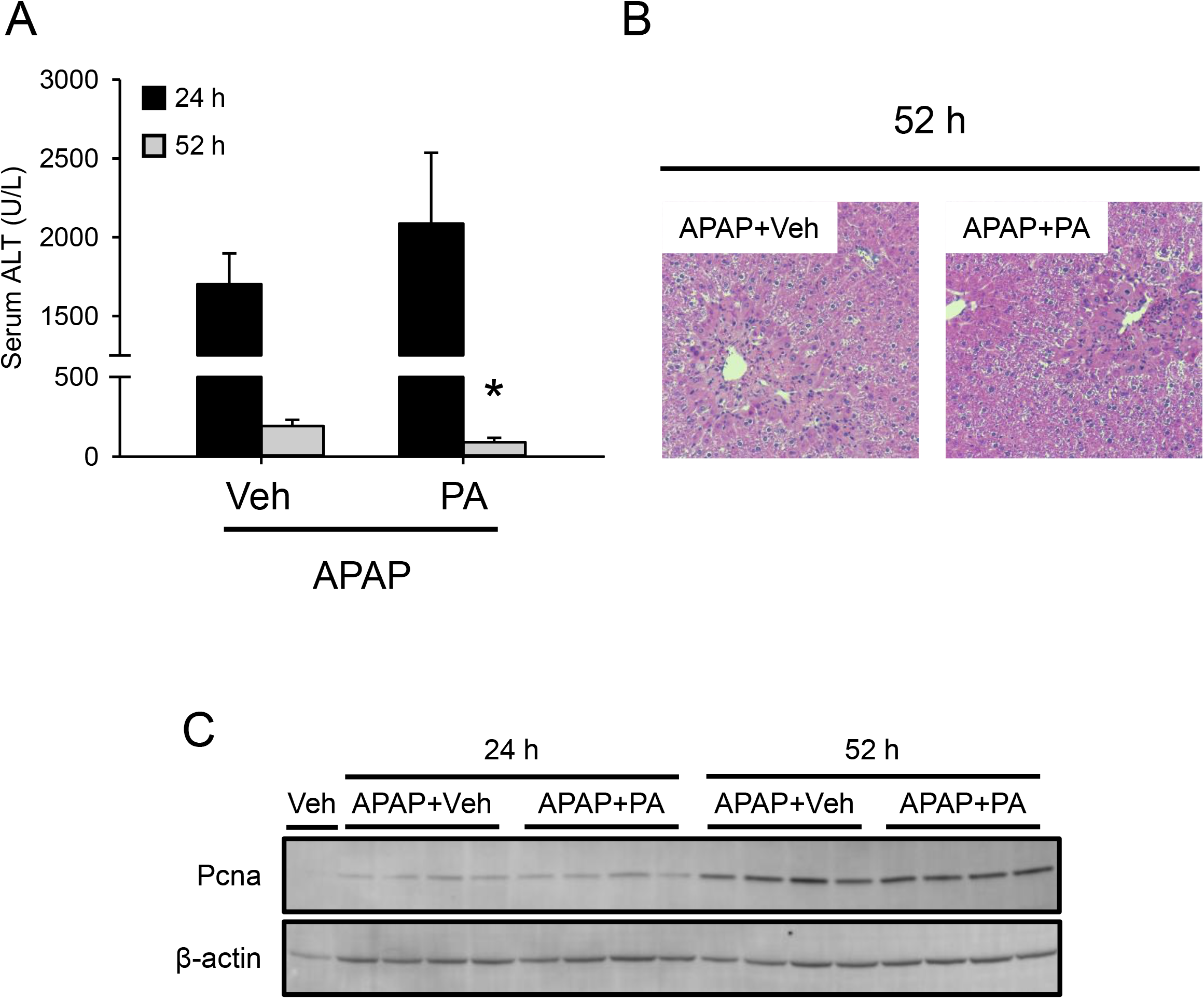
Late post-treatment with exogenous PA does not affect liver regeneration. Mice were treated with 250 mg/kg APAP at 0 h, followed by vehicle (Veh) or PA at 6, 24, and 48 h. Blood and liver tissue were collected at 24 and 52 h. (A) Serum ALT. (B) H&E-stained liver sections. (C) Immunoblot for proliferating cell nuclear antigen (Pcna) and β-actin. Data expressed as mean ± SE for n = 4-5 per group. *p<0.05 vs. APAP plus Veh.

To identify other possible sources of IL-6, we treated mice with PA or vehicle and collected liver, kidney, lung, epididymal white adipose tissue (eWAT), and spleen 4 h later. We chose these tissues because they have high basal IL-6 expression and are known to produce IL-6 in other disease contexts. Interestingly, we observed an 18-fold increase in IL-6 mRNA in eWAT (Fig. 6D). We could not detect differences in the other tissues. These data reveal that adipose tissue is a likely source of increased systemic IL-6 after PA treatment, indicating inter-organ crosstalk between liver and fat.

### 3.5. Exogenous PA does not promote liver regeneration

Finally, because we previously demonstrated that endogenous PA promotes liver regeneration (Lutkewitte et al., 2018; Clemens et al., 2019a) and because IL-6 is a well-known driver of regeneration (Clemens et al., 2019b), we wanted to determine if exogenous PA enhances regeneration and repair after APAP overdose. To test that, we treated mice with APAP at 0 h, followed by exogenous PA or vehicle at 6, 24, and 48 h post-APAP. We selected these late post-treatment time points to avoid an effect on the early injury, which would have decreased liver regeneration secondary to the reduced injury. We then collected blood and liver tissue at 24 and 52 h. Although serum ALT was significantly decreased at 52 h (Fig. 7A), there was no apparent difference in area of necrosis (Fig. 7B) and no change in Pcna (Fig. 7C) between the treatment groups at either time point. These data indicate that exogenous PA, unlike endogenous PA, does not affect liver regeneration after APAP overdose.

## 4. Discussion

Together with our earlier work, the results from this study reveal that endogenous and exogenous PA have different beneficial effects in APAP hepatotoxicity involving different mechanisms of action. We previously demonstrated that endogenous PA accumulates in liver tissue and plasma after APAP overdose in both mice and humans (Lutkewitte et al., 2018). Importantly, inhibition of the PA accumulation had no effect on injury in the mice, but did reduce regeneration and survival by de-regulating GSK3β activity through an effect on Ser9 phosphorylation (Lutkewitte et al., 2018; Clemens et al., 2019a). In the present study, we found that exogenous PA reduces the early injury by increasing systemic IL-6 levels, but has no effect on GSK3β phosphorylation or liver regeneration. These data indicate that exogenous PA or PA derivatives may be a useful adjunct with NAC to treat early APAP hepatotoxicity in patients, but targeting PA-mediated signaling to promote liver regeneration in late presenters will require a different approach.

Our data demonstrate that exogenous PA reduced early injury but had no effect on the major intracellular mechanisms of APAP hepatotoxicity (NAPQI formation, oxidative stress, mitochondrial damage). Transcriptomics analysis then indicated that PA activated IL-6 signaling. Through upstream analysis, we observed activation of IL-6/STAT3 signaling in liver tissue from our PA-treated animals. We then demonstrated that PA does not protect in IL-6 KO mice. Those results are consistent with earlier data demonstrating that systemic administration of exogenous PA dramatically increases circulating levels of IL-6 (Lim et al., 2003). They also confirm the protective role of IL-6 in APAP hepatotoxicity. Masubuchi et al. (2003) reported that IL-6 KO mice have worse injury after APAP overdose. More recently, Gao et al. (2019) observed that administration of exogenous IL-6 is protective. The mechanism by which IL-6 protects is unclear but could involve Hsp70, since Hsp70 and other Hsps are increased in liver tissue after APAP treatment in an IL-6-dependent manner (Masubuchi et al., 2003) and KO of Hsp70 worsens APAP toxicity (Tolson et al., 2006). However, we could not obtain consistent data in support of that hypothesis. Taken together with our data, it seems likely that exogenous PA delays injury through its effects on IL-6 signaling, but some details remain unknown.

Bae et al. (2017) recently demonstrated that exogenously administered lysoPA also protects against APAP hepatotoxicity. PA can be converted to lysoPA by phospholipases, so it is theoretically possible that lysoPA contributed to the protection we observed in our study. However, their data demonstrated that lysoPA protected by 1) preventing early glutathione depletion and increasing glutathione re-synthesis at 6 h post-APAP and by 2) altering JNK and GSK3β activation (Bae et al., 2017), while we could not detect any effect of exogenous PA on either glutathione or kinases in our experiments. These data indicate that PA protected through entirely different mechanisms in our study. However, Bae et al. (2017) also used a 1 h pre-treatment in most of their experiments, which has limited clinical relevance and makes it difficult to directly compare our results.

It is surprising that exogenous PA did not enhance liver regeneration after APAP overdose despite multiple treatments, especially considering the importance of IL-6 in liver repair. IL-6-deficient animals have delayed regeneration after partial hepatectomy, APAP overdose, and CCl4 hepatotoxicity (Cressman et al., 1996; Selzner et al., 1999; James et al., 2003; Rio et al., 2008). On the other hand, Bajt et al. (2003) found that injection of recombinant IL-6 does not enhance regeneration after APAP overdose, and many treatments that do enhance regeneration do not increase IL-6. It may be the case then that basal IL-6 levels are sufficient to aid liver repair, such that reducing IL-6 can blunt regeneration but increasing it has no effect. In any case, IL-6 can clearly influence both early injury and later regeneration in multiple liver disease models, and we need more data to understand the details of those effects.

## 5. Conclusions

Overall, we conclude that post-treatment with exogenous PA likely reduces APAP hepatotoxicity in mice by increasing systemic IL-6, which then activates IL-6 signaling in the liver. Because PA is readily available over-the-counter as a supplement due to its purported ergogenic effects (Shad et al., 2015) and because the combination of PA and NAC protected better than NAC alone in our experiments, exogenous PA or PA derivatives may one day be a useful adjunct with NAC for treatment of early-presenting APAP overdose patients. However, more research is needed to test that possibility. In future studies, we will optimize the dose of PA for protection, test additional treatment regimens and time points, and explore the effects of different acyl chains. We will also test the effects of both endogenous and exogenous PA in other liver disease models.

## Abbreviations

APAP: acetaminophen
ALF: acute liver failure
NAPQI: N-acetyl-p-benzoquinone imine
JNK: c-Jun N-terminal kinase
NAC: N-acetylcysteine
PA: phosphatidic acid
IL-6: interleukin-6
ALT: alanine aminotransferase
GSH: reduced glutathione
GSSG: oxidized glutathione
GSK3β: glycogen synthase kinase 3β
GO:BP: gene ontology:biological processes
Stat3: signal transducer and activator of transcription 3
KC: Kupffer cell
eWAT: epididymal white adipose tissue
Pcna: proliferating cell nuclear antigen
Hif2α: hypoxia-inducible factor
CCl4: carbon tetrachloride
mTORC1: mechanistic target of rapamycin complex 1
LysoPA: lyso-phosphatidic acid
DAG: diacylglycerol

## Acknowledgements

This study was funded in part by a Pinnacle Research Award from the AASLD Foundation (MRM); the Arkansas Biosciences Institute (MRM), which is the major research component of the Arkansas Tobacco Settlement Proceeds Act of 2000; and the National Institutes of Health grants T32 GM106999 (MMC and JHV), R01 DK104735 (BFN), R01 DK117657 (BFN), R42 DK121652 (BFN), R56 DK111735 (BFN), R42 DK079387 (LPJ), and UL1 TR003107 (LPJ, SKM) and TR003108 (LPJ, SKM). We are grateful for expert technical assistance provided by the Dept. of Laboratory Animal Medicine at UAMS (especially Robin Mulkey) and by the Experimental Pathology Core (especially Jennifer D. James, HT(ASCP), HTL, QIHC).

